# Consistency in phonetic categorization predicts successful speech-in-noise perception

**DOI:** 10.1101/2025.09.11.675656

**Authors:** Rose Rizzi, Gavin M. Bidelman

**Affiliations:** Department of Speech, Language and Hearing Sciences, Indiana University, Bloomington, IN, USA; Program in Neuroscience, Indiana University, Bloomington, IN, USA; Cognitive Science Program, Indiana University, Bloomington, IN, USA

**Keywords:** Categorization, speech-in-noise, consistency, gradience

## Abstract

Listeners bin continuous changes in the speech signal into phonetic categories. But they vary in how consistently/discretely they assign speech sounds to categories, which may relate to speech-in-noise (SIN) perception. Yet, it is unclear if and how perceptual gradience, consistency, and other cognitive factors (e.g., working memory) collectively predict SIN performance. Here, we estimated perceptual gradiency and response consistency during vowel labeling and assessed working memory and SIN performance. We found perceptual consistency and working memory were the best predictors of listeners’ composite SIN scores. Our findings emphasize the importance of perceptual consistency over categoricity for noise-degraded speech perception.

## 1. Introduction

To deal with continuous variance in the speech signal, listeners bin speech sounds into phonetic categories ^1,2^. The classic view of categorization suggests that listeners use a small number of auditory cues to inform category membership and discard irrelevant, within-category acoustic information. Yet, studies show listeners have access to both between- and within-category information ^3,4^ and vary in how they weight this information. Some listeners have a more continuous perceptual representation of the speech signal (gradient listener) while others strongly warp the acoustics onto distinct phonetic categories (discrete listener) ^5,6^. A person’s listening strategy, (i.e., degree of perceptual gradiency), can be measured via the slope of their identification curves derived from labeling tokens along an acoustic-phonetic continuum. More gradient listeners have shallower identification ^5^, consistent with weaker categorical percepts and more continuous hearing.

Individual differences in categorization are not restricted only to perceptual gradiency. Listeners also vary in how *consistently* they label tokens along the continuum ^5,7,8^. Consistency may be a more precise measure of individual differences in categorization than slope, accounting for intertrial differences in perceptual reports ^9^. Whether consistency and gradiency are independent measures remains debated ^5,6,8,10^ but measuring both slope (gradiency) and intertrial variance (consistency) allows them to be directly compared as independent factors in speech perception ^11^.

Classic categorization experiments have also used two-alternative forced choice (2AFC) paradigms, requiring listeners to make a binary decision. Consequently, categories might be an artifact of task demands rather than a true perceptual phenomenon ^12,13^. 2AFC tasks confound whether a shallower identification slope represents a more gradient listener or a less consistent labeler ^5,14^. Recent work favors a visual analog scale (VAS) to assess categorization, which allows for more gradiency in perceptual reports ^5,14-16^. Both the slope of the identification curve and intertrial consistency can be quantified and disentangled from VAS responses ^5,14^. For instance, discrete listeners will primarily use the endpoints of the scale to make their reports, but how consistently they choose the same endpoint for a given token may vary. Gradient listeners will have a distribution of responses that moves down the scale mirroring changes in the acoustic cue, though the spread of token-wise distributions may vary (see **Fig. 1A**).

**Figure 1.**
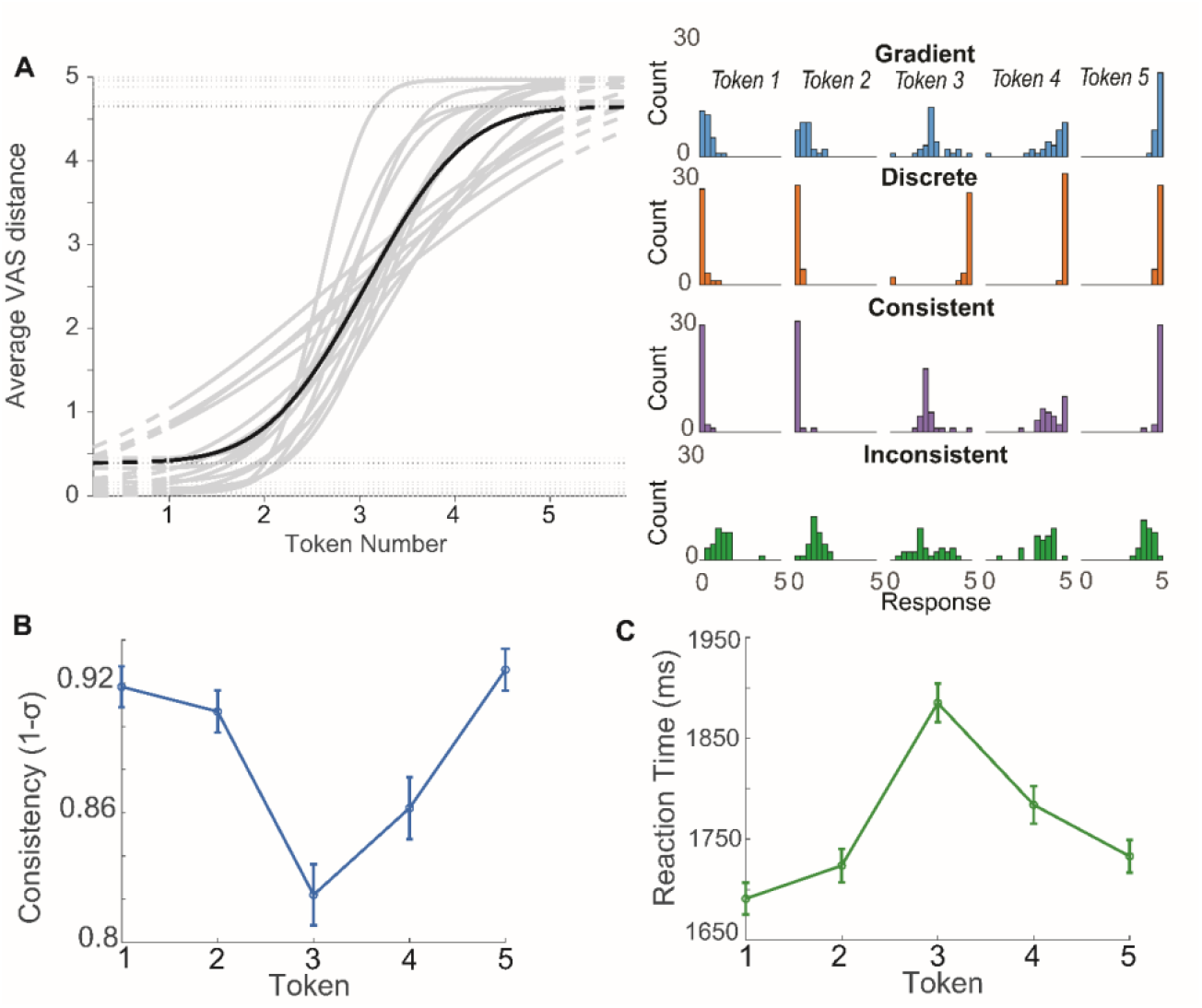
Listeners vary in categorization behavior. (**A**) (*left*) Phoneme identification curves from each participant (gray traces). Grand average (black bold). (*right*): VAS response distributions across continuum steps for representative subjects with different gradiency/consistency response patterns. (**B**) Consistency (1 – σ) across tokens. Listeners’ labeling is less consistent and more variable at the continuum midpoint. (**C**) Reaction times (RTs) for phoneme labeling. Both consistency and RTs are poorer near the continuum midpoint (Tk 3). Error bars = ± s.e.m.

Gradiency in phoneme categorization has been linked to other perceptual processes as well, including increased perceptual flexibility ^5,17^, linguistic diversity ^8^, and indexical variation ^12^. Poorer consistency is also linked to speech-language problems including developmental language disorder and dyslexia ^18^, emphasizing the importance of stable perceptual sound representations to broader auditory-linguistic skills. More consistent listeners also seem to have an advantage for second language learning ^19,20^.

Having a more consistent internal representation of speech sounds might also allow listeners to better parse speech from noise. Maintaining perceptual flexibility with gradient hearing may allow better recovery from ambiguity introduced by noise degradation. However, whether categorization skills predict SIN performance is equivocal. Recent work demonstrates increased consistency and gradience in categorization corresponds with better SIN comprehension ^6,10^. However, others have found this relationship dissolves when controlling for working memory (WM) ^5^, a known cognitive factor important for SIN outcomes ^21-24^. How perceptual (gradiency, consistency) and cognitive (WM) factors contribute to and potentially interact in SIN performance remains an open question. We directly pitted these factors against one another in the current study to assess the degree to which categorization (gradiency, consistency) and WM explain SIN outcomes.

## 2. Methods & Materials

### 2.1 Participants

The sample included *N*=16 normal hearing (pure-tone thresholds ≤25 dB HL; 250-8,000 Hz) adults aged 18-35 (mean = 23.88 ± 4 years; 9 female, 7 male). Participants were right-handed (75% ± 27% laterality) ^25^, had 8.29 ± 6.65 years of music training, and were native speakers of American English. All provided written informed consent in accordance with the Indiana University IRB.

### 2.2 Stimuli and tasks

#### Phoneme categorization

Stimuli consisted of 5 vowel tokens equidistantly sampled from a first formant frequency (F1) continuum changing from /u/ (430 Hz) to /a/ (730 Hz) ^26,27^. Tokens were otherwise identical with respect to F0 (150 Hz), F2 (1090 Hz), and F3 (2350) and were 100 ms in duration (10 ms ramps). Stimuli were presented at 75 dB SPL via PsychoPy (v2023.1.1) ^28^ through Sennheiser HD 280 circumaural headphones. Listeners heard 30 presentations of each vowel (total=150 trials). The interstimulus interval between trials was 500 ms. On each trial, they clicked along a visual analog scale (VAS) with endpoints labelled “oo” and “ah” to report what they heard ^5,6^. Average VAS distance reported and reaction times (RTs) were scored per token.

#### Speech-in-noise (SIN)

We assessed SIN abilities using the QuickSIN ^29^, Hearing in Noise Test (HINT) ^30^, and Words in Noise (WIN) test ^31^. SIN tests were administered via MATLAB at 70 dB HL (QuickSIN), 65 dB-A (HINT), and 75 dB SPL (WIN). The QuickSIN uses 6 low-context sentences with 4-talker babble. The signal-to-noise ratio (SNR) decreases from 25 dB to 0 dB in 5 dB steps with each sentence. The HINT uses 20 higher-context sentences with a speech-shaped noise masker and an adaptive SNR that tracks to the speech reception threshold. For WIN, monosyllabic words follow a carrier phrase “say the word” with six-talker babble at an SNR that decreases by 4 dB every 5 words. We used 35 words for each list. All three SIN assessments calculate the listener’s SNR-50, the SNR at which they can correctly repeat 50% of the keywords.

Lower scores correspond to higher tolerance for noise and thus better SIN performance. For all tests, performance was averaged across two lists. Scores from each of the three SIN tests were z-scored and averaged to arrive at a composite SIN score for each listener. Composite scores were used instead of individual SIN tests to ensure (i) broader generalizability of the results and (ii) putative categorization-SIN relations were not idiosyncratic to a particular SIN measure.

#### Working memory (WM)

We assessed auditory WM capacity using the forward and backward digit span ^32^. A series of digits was verbally presented to listeners (∼1/sec) which varied in sequence length. The number of items progressively increased from 2 to 9 in the forward condition and 2 to 8 in the backward condition (2 trials of each item). Participants scored one point for each sequence correctly recalled. Participants were required to recall the sequence in serial order for the forward condition and reverse order from the presentation for the backwards condition. The test was discontinued if the participant missed both trials. WM spans were computed as the number of correct responses out of 16 and 14 possible points for forward and backwards directions, respectively.

### 2.3 Data analysis

Phoneme identification curves were fit with a sigmoid P = 1/[1 + e^−β1(x−β0)^] and *β1* slopes were estimated using the MATLAB *psignifit* toolbox ^33^. Response consistency of labeling was calculated as 1 – *σ*^*Tk*^, where *σ* is the standard deviation of VAS responses to each token (*Tk*) for each participant ^5^. Reaction times (RTs) in labeling speed were computed as the trimmed mean response latency for each token per condition; RTs outside of 250-3000 ms were deemed lapses of attention or fast guesses and were excluded from analysis ^27,34^.

We used linear mixed-models in R version 4.5 – lme4 package ^35^ to analyze the dependent variables. Separate models were built to test differences in consistency and RTs, with a fixed effect for token (5 levels; Tk1-5) and random intercepts for subject [e.g., RT ∼ Tk + (1|sub)]. Pairwise comparisons were Tukey-adjusted. Effect sizes were calculated as 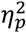. We used partial correlations and linear regression to assess relationships between composite SIN scores, WM digit span scores, and phoneme labeling slope and consistency (SIN ∼ consistency*slope + forward digit + reverse digit). Two participants had missing digit span scores and one with missing HINT and WIN scores. Their data are included in labeling RT and consistency analysis but were treated as missing values for correlations.

## 3. Results

We first confirmed there were individual differences in speech categorization by visualizing identification curves and VAS response distributions. **Fig.1A** shows phoneme labeling functions derived from listeners’ VAS distributions. The steepness of the functions varies across listeners, suggesting differences in the degree of continuous vs. categorical perception (i.e., listening strategy). Representative discrete, gradient, consistent, and inconsistent VAS response patterns across continuum steps are shown on the right. Categorization gradiency is evident in how the responses track across steps, whereas response consistency is evident in the width of distributions around each token. Gradient listeners parametrically shift their perceptual response along the scale with each continuum step, while discrete listeners heavily use endpoints of the scale to respond. More consistent listeners have a smaller spread in their VAS response distributions than inconsistent listeners. Slope and average consistency were weakly correlated [*r*(14) = 0.51, *p* = 0.045], suggesting these measures are partially separable.

We next assessed the effects of stimulus token on labeling consistency. We found a main effect of stimulus token on labeling consistency 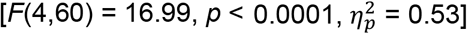. This was driven by less consistent labeling of the ambiguous midpoint token [all (Tk1,2,5) vs. Tk3: *p* < 0.0001, except Tk3-4 contrast: *p* = 0.07] (**Fig. 1B**). Labeling was less consistent for phonetically ambiguous tokens than for endpoint phonetic tokens. Likewise, RTs showed a main effect of token on labeling speeds 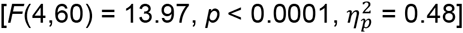, driven by slower RTs for Tk3 [(Tk1,2,4,5) vs. Tk3: *p* < 0.007] (**Fig. 1C**).

We next assessed whether phoneme labeling consistency, slope, and/or WM digit span scores predicted composite SIN scores. Linear regression showed consistency (collapsed across tokens) 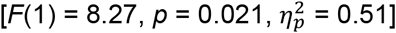 and reverse digit span score 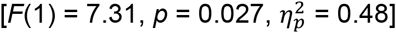 independently predicted SIN scores, while slope 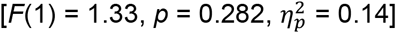, forward digit span 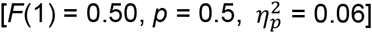, and slope*consistency interaction 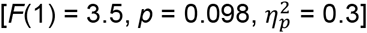 did not. To determine if WM mediated the phoneme consistency-SIN relation, we performed a partial correlation [*ppcor* package in R ^36^] between composite SIN scores and the significant predictors of our model (average consistency and reverse digit span scores – **Fig. 2**). Partialing out the other variables, SIN was still negatively correlated with reverse digit span [*r*(14) = -0.6, *p* = 0.030] and remained marginally correlated with average consistency [*r*(14) = -0.54, *p* = 0.058]. Reverse digit span and consistency were not correlated [*r*(14) = -0.20, *p* = 0.520]. We also found no relationship between WM (reverse digit span) and categorization slope [Pearson’s-*r*: *r*(14) = -0.27, *p* = 0.360].

**Figure 2.**
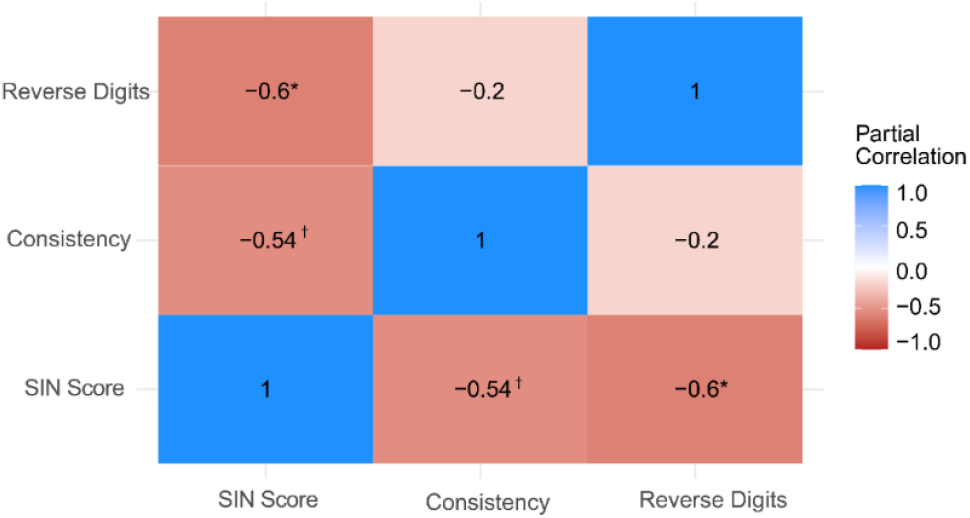
Working memory (reverse digit span) is negatively correlated with SIN perception (composite score). Cell values reflect Pearson correlations between pairwise variables partialled for the remaining third variable. Listeners with better working memory (i.e., larger capacity) perform better on SIN assessments (i.e., lower thresholds). Consistency in phoneme labeling is marginally correlated with SIN. Behavioral slopes in phoneme labeling are not correlated with SIN scores. * *p<*0.05, ^†^*p*= 0.058.

## 4. Discussion

Prior work has implied a potential link between categorization skills (i.e., acoustic-to-phonetic mapping of phonemes) and noise-degraded speech perception ^5,6,10,37^. Here, we explicitly assessed whether phoneme labeling and WM predicted SIN scores measured in the same listeners. We found perceptual labeling was least consistent for more ambiguous phonemes. Importantly, WM capacity and phoneme labeling consistency predicted SIN performance with greater WM and higher consistency corresponding with better (lower) composite SIN scores. These results establish a link between categorization consistency and SIN perception skills, which is not explained solely by cognitive factors.

In VAS phoneme labeling tasks, listeners’ responses were least consistent for midpoint tokens along an acoustic phonetic continuum (Tk3). This aligns with prior work demonstrating slower labeling responses near the continuum midpoint where category membership is ambiguous ^3,6,38^. In speech categorization, tokens lying at a category boundary may be assigned to either category with equal probability, leading to the prototypical sigmoid-shaped identification function (e.g., ^1,2,39^). Using a visual analog scaling (VAS) paradigm, we find more phonetic ambiguity results in less consistent labeling. Responses to tokens with ambiguous category membership are distributed more broadly across the response scale compared to tokens belonging to a clear phonetic category. Consistency is not straightforwardly measured from a conventional 2AFC task ^5,12,14^. However, when measured from VAS paradigm, categorization consistency mirrors RTs and discrimination patterns from 2AFC experiments. If categories were purely an artifact of the 2AFC task ^12^, consistency and RTs from the VAS task should not follow categorical patterns. Our results challenge the notion that categories are an artificial by-product of a 2AFC task structure which forces categorical responses, as tokens with clear category membership are labelled faster and more consistently than those at a phonetic boundary, even in a VAS task which allows continuous responses and promotes gradience.

Labeling consistency and identification curve slope also varied across listeners and were only weakly correlated. More discrete listeners (i.e., with steeper slopes) tended to be more consistent labelers. Whether categorization gradiency and consistency in speech perception are independent constructs is debated ^5,6,8,10^. Instead, listeners might fall into at least one of three sub-types of “speech categorizer” in terms of their perceptual gradiency and consistency: canonically gradient, canonically categorical, or categorical but noisy ^8^. Instead of grouping listeners as in prior work, we treated categorization slope and consistency as continuous measures, allowing for their interaction in our analysis. This revealed consistency in listeners’ perceptual labeling was related to their SIN performance, even after controlling for the degree of gradiency/discreteness in their hearing. While some studies suggest perceptual gradiency is advantageous for SIN recognition ^6,10^, our findings here suggest increased perceptual *consistency* better predicts noise-degraded listening skills.

Aside from perceptual categorization abilities, cognition is also critical for SIN listening. Our results converge with Kapnoula, Winn [5] who similarly found links between categorization and SIN perception were no longer significant when accounting for listeners’ WM. Indeed, after accounting for span capacity, categorization slopes no longer predicted composite SIN scores. Both slope and consistency were not independently correlated with WM, suggesting these categorization measures do not directly depend on WM and may be independent of cognitive abilities. Our finding that consistency was most related to SIN performance aligns with converging work suggesting response consistency is a stable, trait-like property of behavior, and thus more likely to play a role in other perceptual processes ^9,40^. In contrast, perceptual gradiency (as indexed by identification slopes) seems more malleable and may change across different phonetic contrasts and acoustic-phonetic stimuli ^10,17,20^. This questions the generalizability of slope measures as a stable metric of speech categorization abilities (cf. ^37^). Instead, consistency may be a more robust index of perception. In our previous work, we measured categorization “listening strategy” via identification curve slopes ^6^. Future work should consider analyzing gradiency (slope) and consistency as covarying measures of listening strategy for a more complete picture of perceptual abilities.

Expectedly, WM capacity predicted SIN performance. These findings underscore the well-known advantage of superior WM in SIN comprehension ^21-24^. Categorization gradiency is generally independent of cognitive processes like control, attention, or inhibition ^16^. Prior work has found only a weak relationship between gradiency and WM ^5^. Here, we found WM was not correlated with phoneme labeling consistency or gradiency, suggesting these speech perception factors are independent of cognitive abilities. Collectively, our results suggest that perceptual consistency in speech sound labelling may play an underappreciated role in governing SIN comprehension skills, even in normal hearing adults. WM capacity is often difficult to train and thus might be a limited target for programs designed to improve SIN listening ^41^. Our data imply that improving perceptual consistency during speech listening may offer an easier and potentially more robust transfer to SIN outcomes than attempting to train cognitive factors.

## Acknowledgements

Work supported by the National Institute on Deafness and Other Communication Disorders (R01DC016267). Requests for data and materials should be directed to G.M.B.

## Author Declarations

### Ethics Approval

This protocol was approved by the Indiana University Institutional Review Board (23256). Written informed consent was obtained from all participants.

### Data Availability

The data that support the findings of this study are available from the corresponding author upon reasonable request.

